# Exposure to constant artificial light alters honey bee sleep rhythms and disrupts sleep

**DOI:** 10.1101/2023.07.03.547605

**Authors:** Ashley Y. Kim, Aura Velazquez, Belen Saavedra, Benjamin Smarr, James C. Nieh

## Abstract

Artificial light at night (ALAN) is known to create changes in animal behavior in some invertebrates and vertebrates and can result in decreased fitness. ALAN effects have not been studied in European honey bees (*Apis mellifera*), an important pollinator. Colonies can be exposed to ALAN in swarm clusters, when bees cluster outside the nest on hot days and evenings, and, in limited cases, when they build nests in the open. Forager bees maintained in incubated cages were subjected to constant light or dark and observed with infrared cameras. The bees maintained a regular sleep pattern for three days but showed a shift on the fourth day in the presence of continuous light. Bees under constant light demonstrated a 24.05-hour rhythm, compared to a 23.12-hour rhythm in the dark. After 95 hours, the light-exposed bees slept significantly less and experienced significantly more disturbances from their peers. They also preferred to sleep in the lower portion of the cages, which had lower light intensity. These findings suggest that ALAN can disrupt honey bees’ sleep patterns, which may have implications for their behavior and overall colony health.

## INTRODUCTION

Insect declines are historically thought to be driven by climate change, habitat loss, introduced species, and agricultural pesticides (Grubisic et al., 2018). Although a neglected stressor, light pollution covers roughly a quarter of the Earth’s surface (Falchi et al., 2016) and disturbs biological rhythms, orientation, and animal mating behavior and reproductive success (McDonnell et al., 2009). As biodiversity continues to decline due to anthropogenic disturbances, understanding the effects of light pollution on insect systems is increasingly crucial for conservation. Artificial Light at Night (ALAN) has ecosystem impacts (Falcon et al., 2020) because it can interfere with pollination networks and disrupt plant fitness and food webs (Knop et al., 2017). For example, ALAN can significantly decrease the amount of time that moths spend feeding, which could contribute to their decline and negatively impact the pollination services they provide (van Langevelde et al., 2017). However, the underlying effects of ALAN on pollinator behavior are relatively unknown (Falcon et al., 2020), particularly for insects such as social bees in which cooperative foraging is essential for colony fitness. Because honey bees are one of the most important pollinators in natural and agricultural ecosystems (Hung et al., 2018; Khalifa et al., 2021), understanding the effects of ALAN on honey bees is crucial.

In general, honey bees prefer to nest in dark cavities where the only light source would be the hive entrance. As ALAN increases, the nest entrance light source can be constant during a 24 h period. When honey bees travel to a dark part of the nest, the sensitivity of light in their eyes increases by 1000-fold (Wolf & Zerrahn-Wolf, 1935). This photic adaptation may be disrupted when constant light illuminates parts of the hive. In particular, nurse honey bees live inside a normally dark hive and do not show strong circadian rhythms until they become foragers when they are regularly exposed to daylight, an important zeitgeber (Rodriguez-Zas et al., 2012).

There are multiple ways in which honey bees can be exposed to ALAN if they are outside their enclosed nests and close to ALAN. Honey bees can cluster outside the nest during extreme heat, a phenomenon known as “bee bearding”, typically during hot (≥38 °C) and humid nights and days that occur from mid-June to August in areas such as Europe (Hamdan, 2015). This response to extreme heat stress is part of colony thermoregulation (Stabentheiner et al., 2021), and may occur more frequently as global temperatures increase (Thomas, 2010). Honey bee colonies also reproduce through swarming, a process in which honey bee colonies split into two or more colonies as a means of reproduction. The swarming process typically takes hours to complete but can also span multiple days (Seeley, 2003). The frequency of swarms in urban environments with ALAN has been observed to increase when winter precipitation increases (Baum et al., 2008). Finally, bees can be exposed to ALAN when they build nests in the open, not inside cavities. Although *A. mellifera* is a cavity-nesting species, beekeepers and others have reported cases in temperate climates in which colonies have not nested inside a cavity but, instead, in the open where they can be exposed to ALAN.

Honey bee circadian rhythms are free running under both constant light or dark conditions, can be entrained by light-dark (LD) cycles, can be phase-shifted by pulses of light, and can vary from 20-26 h cycles (Moore, 2001). These rhythms also help regulate sleep, which can be separated by stages of light and deep sleep, and total antennal immobility is thought to be the deepest state of sleep (Kaiser, 1988). Sleep in honey bees is a dynamic process that consists of relative immobility, discontinuous ventilation, and an increased response threshold to external stimuli. Though generally immobile, legs, tarsi, or antennae can twitch, and the body can exhibit subtle movements caused by nestmates’ interactions. Bees need sleep to effectively communicate with nestmates (Klein et al., 2018). Sleep is therefore important for colony fitness because the honey bee dance language can significantly enhance colony fitness (Sherman and Visscher, 2002).

We hypothesized that foragers collected from colonies will maintain, for some time, their natural circadian rhythms, as measured in bee sleep, under the normally dark conditions inside the nest, but would have these sleep rhythms disrupted when they are exposed to constant illumination. We also hypothesized that bee stress would be significantly higher under constant light conditions due to disruptions in sleep behavior.

## METHODS

### Constant illumination harmed honey bee sleep

In Spring 2021, three trials (*n*=6 colonies) were conducted to observe sleep behavior under both dark and light treatments. We used six different colonies. In total, we tested 120 bees, each cage ranging from 14-28 individuals. In La Jolla, California, we captured *Apis mellifera ligustica* foragers from sucrose solution feeders (2.5 M) at nest entrances and transported the bees from the UCSD Biology Field Station (BFS) to the lab. Foragers had therefore been exposed to natural cycles of light and dark for multiple days before their capture. Honey bee foragers were placed into two different experimental cages. The experimental cages were made from transparent acrylic (12 cm long × 8 cm wide × 12 cm high) and had a sliding door on the top with holes for ventilation with another hole to insert a 5 mL syringe containing 2.5M sucrose solution. We also placed a half strip of a queen mandibular pheromone (QMP, Temp Queen Pheromone, catalog number DC-705, Meyer Bees, Chicago, USA) because queen presence, which can be simulated by QMP, is required for normal worker vitality (Kaminski et al., 1990). The experimental cages were placed inside an incubator (24 °C, 50% relative humidity) and bee behaviors were recorded with two infrared-sensitive Lorex 4K cameras (LNE894AB) connected to a digital recording system (Lorex N842, Lorex USA) for 24 h for five consecutive days. Moderate-intensity infrared LED lights (850 nm), which are invisible to bees because they cannot typically detect light wavelengths higher than 550 nm (Chittka et al. 1997) were used to illuminate the bees in both cages for video recordings.

To emulate constant illumination from buildings and streetlights, white LED light bulbs were used with a brightness of 2.7±0.1 μmol/m^2^/s (measured inside at the level of the cage bottom (McMunn et al., 2019). We placed the light treatment cage in a light-proof box made of cardboard and aluminum foil and had an exposed LED bulb (light treatment). The dark treatment cage was placed into the incubator and had no exposure to constant light. We measured photon flux with an ePAR light sensor (Apogee Instruments, Model DLI-600, spectral range of 390-760 nm) placed inside the cage and flush with the cage top and, separately, on the cage floor by cutting a circular hole into an identical cage in which we inserted the top of the light sensory, level with the cage floor and pointed up at the cage top through which light entered. We made 15 measurements of photon flux at the top and 15 measurements of photon flux at the bottom of the cage.

We did not expose bees to a third condition, 12 h of light and 12 h of dark each 24 h, because we wished to simulate the effects of foragers continuously exposed to light for five days, as could occur when colonies form bee beards (Hamdan, 2015) or swarm or nest in the open near artificial light or exposed to dark for five days. Honey bees may remain in dark conditions for prolonged periods, either as a result of overwintering inside the colony (Winston, 1987) or in response to poor weather (Clarke and Robert, 2018). We, therefore, simulated a situation in which bees were kept in the constant dark for five days (our control, dark situation).

To score the sleep data, we looked at the first five minutes of each hour to count how many foragers were sleeping (sleep markers: antennal immobility, non-pulsating abdomen, and leg immobility) and the number of foragers alive or dead on each recorded day. We chose a five-minute time interval that included our sleep markers based on previous literature methodology observing honey bee sleep (Eban-Rothschild et al., 2008) and fruit fly sleep (Tomita et al., 2017). This interval is used as a marker that a honey bee is in a sleep stage. Observers were trained to count forager bees sleeping on the comb and the sucrose syringe. We defined bee sleep as antennal immobility and remaining completely stationary for the entire five-minute observation period.

### Potential behavioral stressors

Based upon visual inspection of the data, we detected a clear divergence in sleep between bees experiencing dark vs. light treatments after 90 h. We, therefore, analyzed the video recordings after this time point for evidence of light avoidance or sleep disruption. Observers who had carefully watched and scored a total of 46.3 h of video (first five minutes of each hour within a 24 h period) for bee sleep noted three potential markers of bee stress that differed between the dark and light treatments: (1) bees sleeping on the bottom of the cage, where light levels were significantly lower (see Results) or (2) disturbance of a sleeping bee by a moving nestmate. Within the first 5 min of each hour, observers therefore (1) scored if the bee was sleeping in the upper or lower 50% of the cage, and (2) counted the number of times each sleeping bee (see definition above) was moved by physical contact with a non-sleeping bee passing by. To distinguish between bees that just happened to wake up and move when another bee passed by from a bee that was sleeping and disturbed, we only scored a sleep disturbance if the sleeping bee settled back into an immobile state immediately after being disturbed.

### Statistics

We used JMP Statistical Software v16.0.0 and built Repeated-Measures Mixed Models (REML algorithm) to examine the effects of treatment, cumulative time since the start of the trial (both fixed effects), and their interaction (with cage identity a random effect nested within treatment) on the proportion of sleeping bees.

To compare *the light levels* in the upper and lower part of the cage, we used a *t*-test comparing light levels in both cage sections. To analyze the effects of potential *behavioral markers of stress*, we used Repeated-Measures Mixed Models (REML algorithm) with cage ID nested within treatment (light or dark) as a random effect, cumulative time from the start of the experiment as the time effect (fixed effect), and the interaction of cumulative time x treatment (fixed effect). We used *survival analysis* to determine the effect of treatment on bee survival over the entire trial.

Due to the loss of individuals over time, we used *N*=1 hypothesis generation approaches to test consistency with the hypothesis that constant light and constant dark would have differing impacts on sleep rhythmicity (Smarr et al. 2015 and De Groot et al. 2017). These analyses were carried out in Matlab 2022b (The Mathworks Inc., Natick, MA, USA). To assess the maintenance of sleep rhythmicity within this framework we took two approaches: 1) comparing fit to sine waves, and 2) comparing phase distribution per cycle.

For sine fitting, sine waves were generated by fitting to the data per group divided into two-day intervals (i.e., data in each group *z*-scored by the first two days, and separately *z*-scored by the next two days). We did not use data from the last day because there were too few bees remaining alive for these rhythmicity analyses. The sine fit was then evaluated with a Pearson correlation between the sine and the fit data.

For phase distribution analysis, “days” were calculated as 2 π radian oscillations, to account for the different free-running periods of each nest under constant conditions. The phase distribution agreement was calculated by plotting the proportion of sleeping individuals by the phase of each daily cycle, as opposed to by clock time. Cycles were defined by taking the trough midpoint of each 24 h interval per nest as a phase marker and then re-interpolating the data to redistribute them between troughs over 2 π radians. In this way, data distributed across multiple cycles with different periods (all roughly but not precisely 24 h) could be compared precisely as distributed across 2 π radians. Data were then plotted radially as a mean of nests per condition over phase, and the area of overlap was calculated for the first two and the next two days per condition. We also calculated the proportion of area from each condition within the 1 π radian centered on the trough, and the 1 π radian opposed.

## RESULTS

### Sleep

There was no significant effect of treatment (*F*_1,4_=0.13, *P*=0.74), but a significant effect of cumulative time (*F*_1,459_=9.15, *P*=0.003) and the interaction of treatment x cumulative time (*F*_1,459_=10.25, *P*=0.002); the proportion of bees in the light treatment, but not the dark treatment, tended to increasingly sleep more throughout daylight hours as time progressed (**Fig. 1A**). This analysis focused on differences between the treatments, but not on the inherent circadian periodicity. Around Day 4 (95 h after the start of the trial), there was a crossover point between the two treatments. There was a significant effect of treatment (*F*_1,34_, *P*=0.0046), time (*F*_1,4_ = 7.25, *P*=0.0011), and their interaction (*F*_1,34_=6.68, *P*=0.014). Bees exposed to constant light slept significantly less than bees in the dark treatment (**Fig. 1A**).

**Figure 1.**
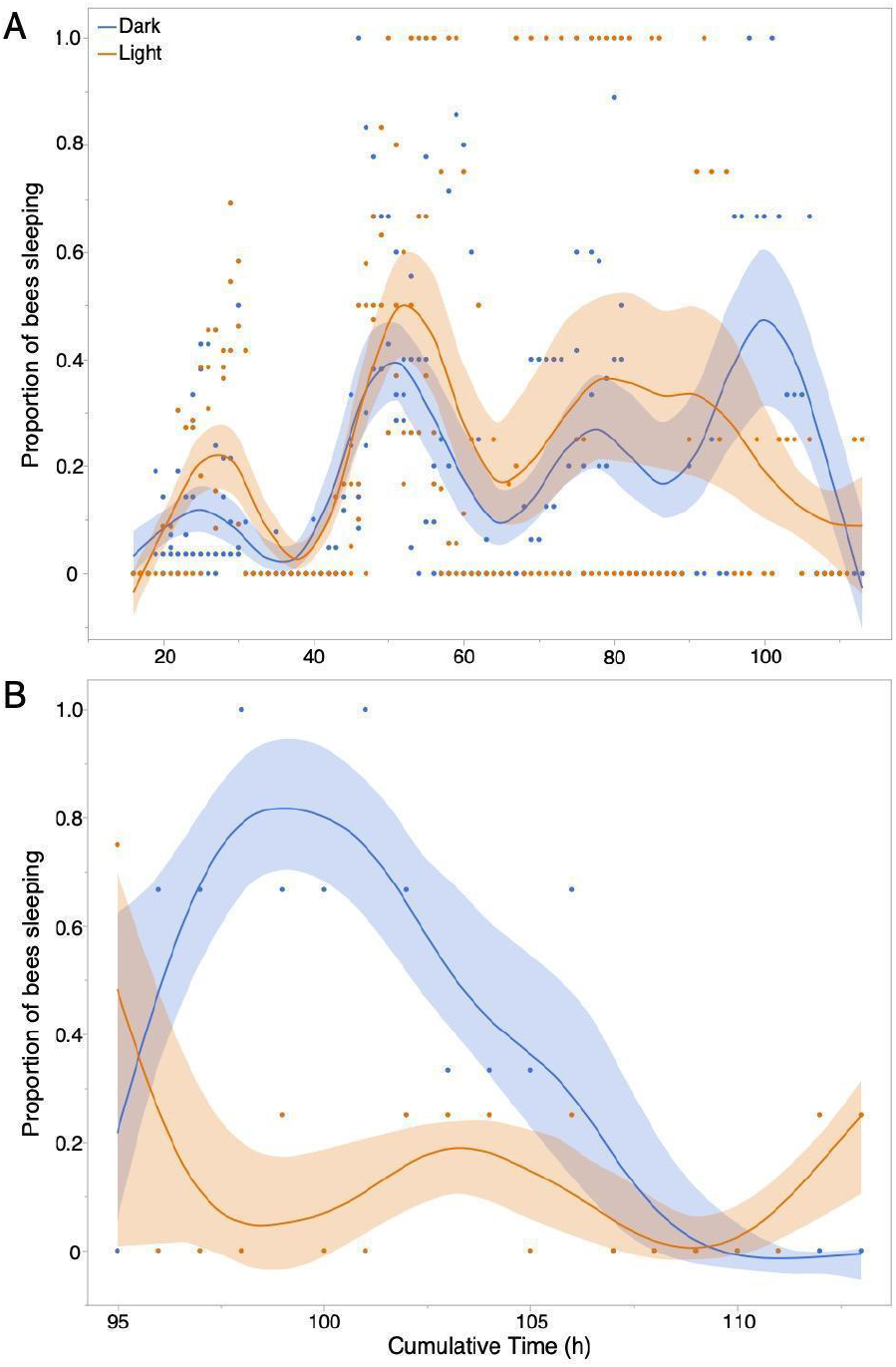
Bee sleep follows a roughly circadian pattern over the entire experiment, but changes after about 90 h in bees under constant illumination. A) Over the entire experiment, bee sleep changed significantly over time (*P*=0.0002). B) From 95 h until the end of the experiment, there was a significant effect of treatment on sleep (*P*=0.005) and a significant effect of time (*P*=0.001). The significant interaction of treatment x time (*P*=0.014) is shown in the different trend lines. Bees exposed to constant light slept less than bees kept in the dark, the normal state inside the nest. All data points are shown, along with best-fit spline curves and 95% confidence intervals.

There was no significant difference between the survival of bees experiencing either treatment (Log-Rank Chi-square = 1.72, 1 d.f., *P*=0.19). The median survival time was 53 h.

### Sleep rhythmicity

Sine fitting of bee sleep under the light and dark treatments revealed comparable fits in the first two days for each condition (first two days: light: *r* = 0.81; dark: *r* = 0.80). By contrast, a sine fit for the next two days revealed a substantial decline in the light but not the dark treatment (last two days: light: *r* = 0.29; dark: *r* = 0.83, **Fig 2A**).

**Figure 2.**
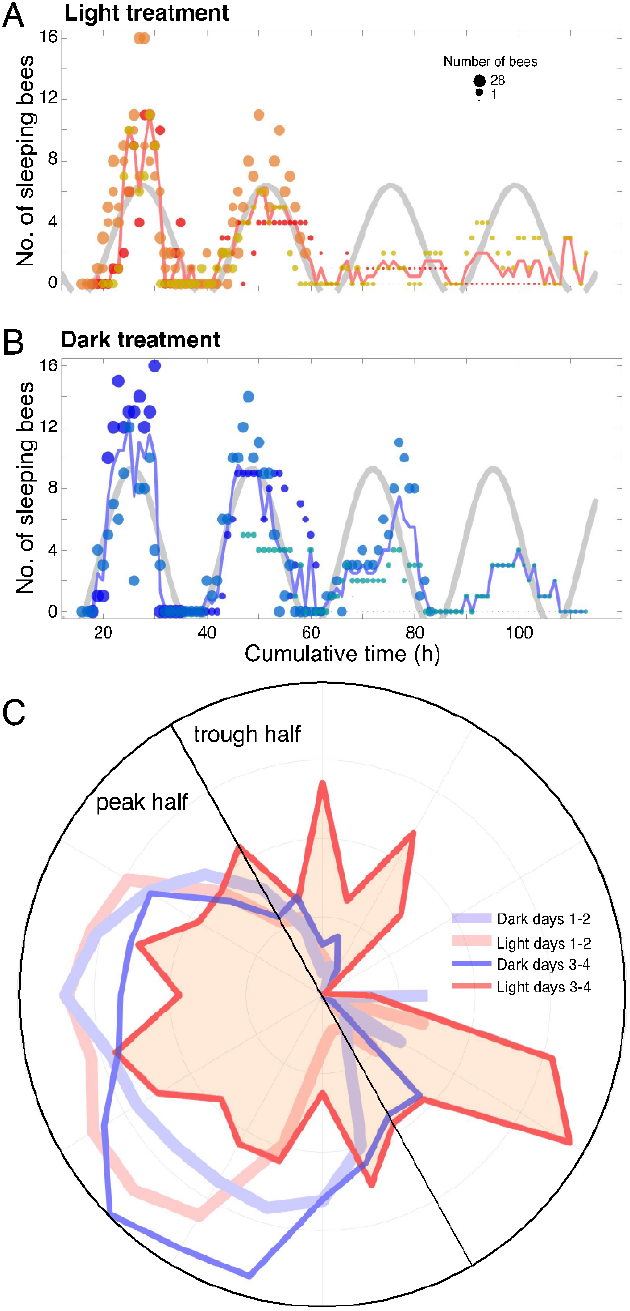
The effects of light vs. dark exposure on bee sleep rhythmicity over time. The sine fit declines under the A) light treatment but not the B) dark treatment. The sine wave was fit to the z-score for the first 2 days and is shown in both panels for clarity, while the fit was evaluated on the last two days separately, as described in Methods. Dot colors represent individual nests by condition, and dot size is proportional to the number of individuals remaining alive. C) Radial graph of the number of sleeping individuals (the number of sleeping individuals is proportionate to the radius for the first two days of each treatment and the second two days of each treatment. The second two days of light exposure are filled in to highlight its difference from the other three groups. “Trough half” is defined as the 1 π radians centered on the daily trough, while “peak half” is defined as the opposing 1 π radian. Units of the area are given in arbitrary units (AU).

Phase distribution analysis revealed a similar finding: the proportion of bees sleeping in the light treatment is substantially divergent between the initial two days as compared to the next two days to the area of overlap (see Methods: Statistics) of bees in the dark from the first two days with the next two days (light: 195 Arbitrary Units (AU) vs dark: 395 AU). This loss of overlap was due to the redistribution of sleep across time (phase) specifically in the last two days of continuous light only (**Fig. 2B**). This is illustrated by the proportion of area under the curve in the 1 π (180 degrees) centered on the daily trough as compared with the proportion of area under the curve within the opposing 1 π. On days 3-4, for bees exposed to continuous light, only 55.7% was in the peak half, whereas for all other conditions (dark 1-2 days, dark 3-4 days, and light 1-2 days bees), 93.7-94.6% of the area was in the peak half (**Fig. 2C**)

### Potential behavioral stressors

There was 7-fold more light in the upper as compared to the lower part of the cage (upper part of the cage: 18.2±0.1 μmol/m^2^/s; lower part of the cage 2.7±0.1 μmol/m^2^/s; t-test, *t*_28_=315.27, *P*<0.0001). In the light treatment, bees slept significantly more often in the lower part of the cage where there was less light. Bees in the dark treatment slept higher up in their cages as compared to bees in the light treatment (treatment effect: *F*_1,34_ = 19.62, *P*<0.001, **Fig. 3A**). Time and the interaction of time x treatment were not significant (F ≤0.48, *P* ≥ 0.49).

**Figure 3.**
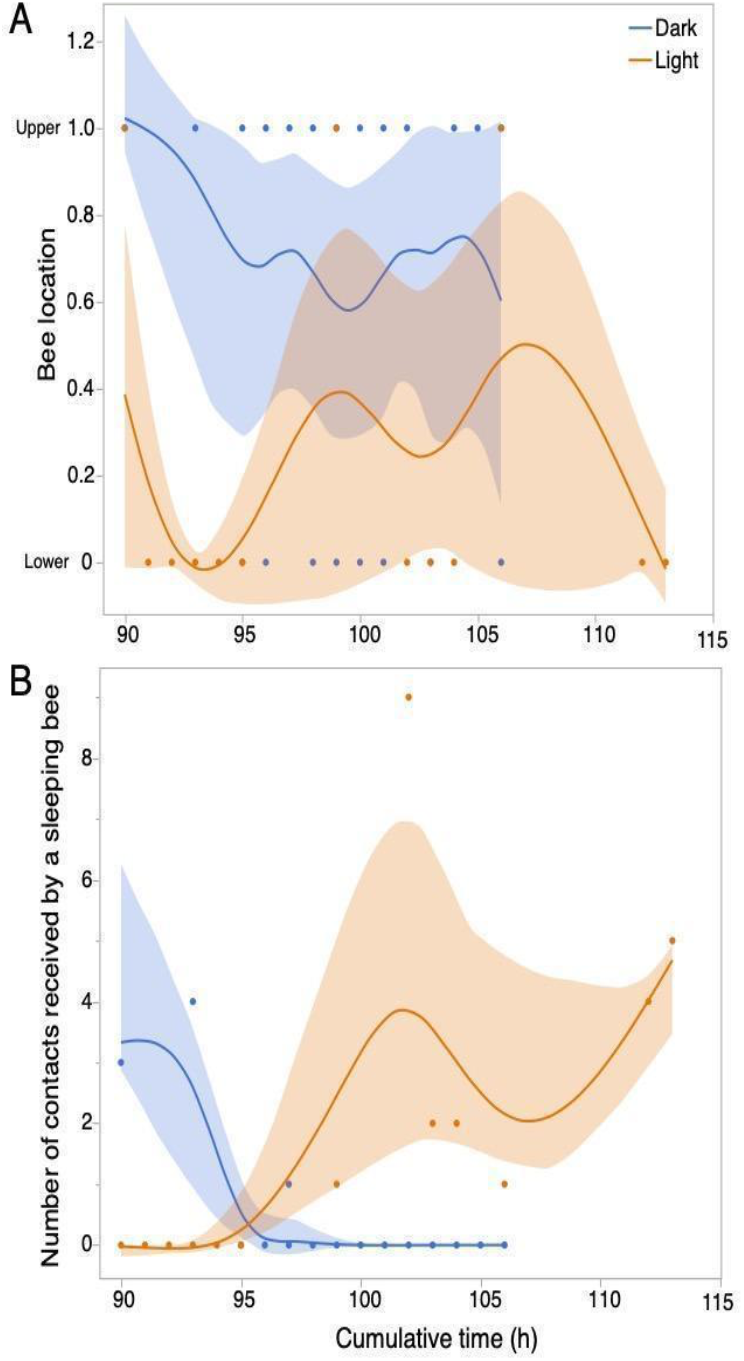
Potential markers of bee stress related to light exposure. A) Bees experiencing constant light slept significantly more often on the bottom of the cage as compared to bees experiencing constant dark at the end of the experiment (≥90 h). There was a significant effect of treatment (*P*<0.001), but no significant effect of time or the interaction of treatment x time (*P* ≥0.49). B) Bees in the light treatment received more potentially disruptive contacts than bees in the dark treatment. There were significant effects of treatment (*P*=0.004) and the interaction of treatment x time (*P*<0.001), which is shown in the different slopes of the spline curves over time (*P*<0.001). All data points are shown, along with best-fit spline curves and 95% confidence intervals.

For physiological markers of stress, bees in the light treatment experienced more disruptive contacts than bees in the dark treatment (*F*_1,44_ = 9.44, *P*=0.004, **Fig. 3B**). Time had no significant effect (*F*_1,44_ = 1.85, *P*=0.18), but there was a significant interaction of time x treatment (*F*_1,34_ =48.53, *P*<0.001) as bees in the light treatment experienced more disruptive contacts. On average, within a 5 min interval, a sleeping bee received 1.09±2.24 contacts from non-sleeping nestmates in the light treatment and 0.31±0.97 contacts from non-sleeping nestmates in the dark treatment (3.5-fold more disruptive contacts in the light as compared to the dark treatment).

## DISCUSSION

In typical dark conditions inside the nest, honey bees demonstrate a stable circadian sleeping rhythm. However, exposure to continuous illumination extends this rhythm to 24.05 hours (compared to 23.12 hours in complete darkness) and appears to cause a loss of rhythmicity after 72 h. Bees in the light treatment slept less after 72 h than their counterparts in the dark treatment. Additionally, continuous light diminished honey bee sleep after roughly 90 hours and prompts behavioral alterations such as increased sleeping in dimmer areas and frequent physical disturbances from non-sleeping bees. This suggests that constant light exposure could be detrimental to bees, leading to sleep loss and reduced sleep quality.

In our experiment, bees were subjected to multiple days of dark or light. In nature, foragers can remain inside their nests for extended periods of time because of poor weather, particularly rainfall or cold, with the most extreme examples occurring in overwintering colonies (Winston, 1987). Thus, it is reasonable to expect that foragers have evolved to cope with extended periods of darkness, and we confirmed that, at least on a scale of five days, forager sleep was not disrupted.

We did not periodically remove dead bees or change out their sucrose solution, as is standard practice with caged bee studies (Williams et al. 2013), to avoid providing a zeitgeber that could alter circadian rhythms. The trials were thus limited to five days. A longer trial period—provided that the experimenters can provide fresh food and remove dead bees without introducing potential time information to synchronize or alter bee circadian rhythms—could provide greater insight into how constant light disrupts bee circadian rhythms. Another limitation of our study is that we ran three trials with six colonies and 120 total bees. A larger number of trials could have allowed us to run more detailed analyses on the effects of constant light on bee sleep rhythms. However, we were still able to identify a clear disruptive effect of light on sleep (*P*=0.0046), trends in changed sleep rhythmicity (**Fig. 2**), where bees slept (*P*<0.001), and the behavioral disruption of sleep (jostling, *P*=0.004). In the dark condition, these contacts were higher before 90 h and then decreased to 0 at 95 h, when the disruptive contacts significantly increased for the constant light bees. This likely occurred because dark condition bees, at this time point, were in a non-sleep phase (**Fig. 1A**), and thus there were very few sleeping bees to be disrupted by other bees. Although there were light treatment bees sleeping during this time period, the movements of other bees that disrupted their sleep appear only to have increased after 95 h. We speculate that the loss of normal circadian sleep rhythms contributed to the behavior of these “sleep disruptor” bees.

Given our findings, future studies may focus on the effects of artificial light on the clock genes expressed in bee brains (Abreu et al., 2018) and how exposure to constant light may alter foraging patterns (Moore, 2001). If foraging efficiency declines, so does the colony fitness for the hive. In addition, the stressful effects of sleep deprivation should be explored. Klein et al. (2010) found that sleep restriction in honey bees significantly lowers their waggle dance precision and thereby lowers the foraging efficiency of signal receivers. Fruit flies like *Drosophila melanogaster* under constant artificial light have been shown to have a significant decrease in fecundity rates and adult survival (McLay et al., 2017). Frequent or repeated exposures to light at night, which could occur with growing light pollution in urban areas and can be exacerbated by bee bearding, the clustering of bees outside the nest on days and nights with higher than normal temperatures, may be chronic stressors that, like other chronic stressors, can lead to changes in animal performance (e.g. growth, endurance, disease resistance) and behavioral patterns (e.g. feeding, aggression, reproduction) (Kay et al., 2021). Studies that have measured different honey bee behavioral and physiological stress responses, including cellular stress responses (Even et al., 2012) may be a useful model for studying different ways in which constant light exposure affects honey bee behavior and physiology. In addition, Sauer et al. (2004) have shown that honey bees can compensate for a sleep deficit by intensifying their sleep. However, the degree to which such sleep compensation can correct for extended sleeplessness induced by constant light remains to be determined.

Studying the effects of ALAN on honey bee swarms in the field, or non-enclosed colonies, would be valuable, particularly in urban settings. In addition, there is an increase in urban beekeeping (Egerer and Ingo 2020). Although this can provide benefits by bolstering pollinator populations, this surge in urban colonies could potentially expose a larger proportion of bee populations the constant artificial light. The urbanization and industrialization of human populations had led to a rise in ailments related to circadian disruption. This urbanization of pollinators presents a significant, worrying, and yet largely unexplored environmental challenge to honey bees. Finally, it may be useful to consider how other honey bee species, like *Apis dorsata* and *Apis florea*, which typically nest in the open, can cope with artificial light at night. Although artificial light has not played a role in the evolutionary history of these species, they are subject to periodic bright moonlight, depending upon their hive locations. Understanding their behavioral and physiological adaptations to light at night could inform our broader understanding of the impact of ALAN on bee populations.

Finally, we should consider how to mitigate the adverse effects of artificial light at night by designing more wildlife-friendly lighting solutions that reduce light pollution. There have been recent strides to develop lighting that minimizes harm to insects and can propose tangible solutions for new urbanization (Deichmann et al., 2021). Incorporating such ecological considerations into our urban planning strategies could potentially safeguard the well-being of multiple pollinators and ensure the sustainability of our urban ecosystems.

## Author contributions

AYK and JCN designed the experiments. AYK conducted the experiments, analyzed the data, and wrote the paper. AV and B Saavedra helped to collect the data. B Smarr and JCN analyzed the data and helped to write the paper. JCN provided research support and funding.

## Data and materials availability

Data will be available upon publication at Zenodo.com at the DOI: 10.5281/zenodo.7992488.

## Acknowledgments

We would like to thank Barrett Klein and the anonymous reviewers for their helpful comments, which have significantly improved this manuscript.

## REFERENCES

Abreu, F. C. P., Freitas, F. C. P. and Simões, Z. L. P. (2018). Circadian clock genes are differentially modulated during the daily cycles and chronological age in the social honeybee (Apis mellifera). Apidologie 49, 71–83.

Baum, K. A., Tchakerian, M. D., Thoenes, S. C., & Coulson, R. N. (2008). Africanized honey bees in urban environments: A spatio-temporal analysis. Landscape and Urban Planning, 85(2), 123–132.

Chittka, L., & Waser, N. M. (1997). Why red flowers are not invisible to bees. Israel Journal of Plant Sciences, 45(2-3), 169–183.

Clarke, D. and Robert, D. (2018). Predictive modelling of honey bee foraging activity using local weather conditions. Apidologie 49, 386–396.

De Groot, M., Drangsholt, M., Martin-Sanchez, F. J. & Wolf, G. (2017). Single Subject (N-of-1) Research Design, Data Processing, and Personal Science. Methods Inf Med 56, 416–418.

Deichmann, J.L., Ampudia Gatty, C., Andía Navarro, J.M., Alonso, A., Linares-Palomino, R. and Longcore, T. (2021). Reducing the blue spectrum of artificial light at night minimises insect attraction in a tropical lowland forest. Insect Conserv Divers, 14: 247–259.

Eban-Rothschild, A. D., & Bloch, G. (2008). Differences in the sleep architecture of forager and young honeybees (Apis mellifera). Journal of Experimental Biology, 211(15).

Egerer, M. and Ingo, K. (2020). “Confronting the modern gordian knot of urban beekeeping.” Trends in Ecology & Evolution 35.11:956–959.

Eubank, R. L. (1999). Nonparametric Regression and Spline Smoothing. 2nd ed. Boca Raton, Florida: CRC.

Even, N., Devaud, J.-M. and Barron, A. B. (2012). General stress responses in the honey bee. Insects 3, 1271–1298.

Falchi, F., Cinzano, P., Duriscoe, D., Kyba, C.C., Elvidge, C.D., Baugh, K., Portnov, B.A., Rybnikova, N.A. and Furgoni, R. (2016). The new world atlas of artificial night sky brightness. Science advances, 2(6), p.e1600377.

Falcon, J., Torriglia, A., Attia, D., Vienot, F., Gronfier, C., Behar-Cohen, F., Martinsons, C. and Hicks, D. (2020). Exposure to Artificial Light at Night and the Consequences for Flora, Fauna, and Ecosystems. Front Neurosci 14, 602796.

Grubisic, M., van Grunsven, R. H. A., Kyba, C. C. M., Manfrin, A. and Hölker, F. (2018). Insect declines and agroecosystems: does light pollution matter? Annals of Applied Biology 173, 180–189.

Hamdan, K. (2015). The Phenomenon of Bees Bearding. Bee World 87, 22–23.

Hung, K.-L. J., Kingston, J. M., Albrecht, M., Holway, D. A. and Kohn, J. R. (2018). The worldwide importance of honey bees as pollinators in natural habitats. Proceedings of the Royal Society B 285, 20172140–8.

Kaiser, W. (1988). Busy bees need rest, too: Behavioural and electromyographical sleep signs in honeybees. J. Comp. Physiol. A. 163, 565–584.

Kaminski, L. A., Slessor, K. N., Winston, M. L., Hay, N. W. and Borden, J. H. (1990). Honeybee response to queen mandibular pheromone in laboratory bioassays. Journal of Chemical Ecology 16, 841–850.

Kay, C. B., Delehanty, D. J., Pradhan, D. S. and Grinath, J. B. (2021). Climate change and wildfire-induced alteration of fight-or-flight behavior. Climate Change Ecology 1.

Khalifa, S. A. M., Elshafiey, E. H., Shetaia, A. A., El-Wahed, A. A. A., Algethami, A. F., Musharraf, S. G., AlAjmi, M. F., Zhao, C., Masry, S. H. D., Abdel-Daim, M. M. et al. (2021). Overview of Bee Pollination and Its Economic Value for Crop Production. Insects 12.

Klein, B. A., Klein, A., Wray, M. K., Mueller, U. G., Seeley, T. D. and Robinson, G. E. (2010). Sleep deprivation impairs precision of waggle dance signaling in honey bees. Proceedings of the National Academy of Sciences of the United States of America 107, 22705–22709.

Klein, B. A., Olzsowy, K. M., Klein, A., Saunders, K. M. and Seeley, T. D. (2008). Caste-dependent sleep of worker honey bees. Journal of Experimental Biology 211, 3028–3040.

Klein, B. A., Vogt, M., Unrein, K., & Reineke, D. M. (2018). Followers of honey bee waggle dancers change their behaviour when dancers are sleep-restricted or perform imprecise dances. Animal Behaviour, 146, 71–77.

Knop, E., Zoller, L., Ryser, R., Gerpe, C., Hörler, M. and Fontaine, C. (2017). Artificial light at night as a new threat to pollination. Nature Publishing Group 548, 206–209.

McDonnell, M. J., Hahs, A. K. and Breuste, J. H. (2009). Ecology of cities and towns: a comparative approach: Cambridge University Press.

McLay, L. K., Green, M. P., & Jones, T. M. (2017). Chronic exposure to dim artificial light at night decreases fecundity and adult survival in Drosophila melanogaster. Journal of Insect Physiology, 100, 15–20.

McMunn, M. S., Yang, L. H., Ansalmo, A., Bucknam, K., Claret, M., Clay, C., Cox, K., Dungey, D. R., Jones, A., Kim, A. Y., et al. (2019). Artificial light increases local predator abundance, predation rates, and herbivory. Environ Entomol 48, 1331–1339.

Moore, D. (2001). Honey bee circadian clocks: behavioral control from individual workers to whole-colony rhythms. Journal of Insect Physiology 47, 843–857.

Rodriguez-Zas, S. L., Southey, B. R., Shemesh, Y., Rubin, E. B., Cohen, M., Robinson, G. E. and Bloch, G. (2012). Microarray analysis of natural socially regulated plasticity in circadian rhythms of honey bees. Journal of Biological Rhythms 27, 12–24.

Sauer, S., Herrmann, E., & Kaiser, W. (2004). Sleep deprivation in honey bees. Journal of Sleep Research, 13(2), 145–152.

Seeley, T.D. (2003) Consensus building during nest-site selection in honey bee swarms: the expiration of dissent. Behav Ecol Sociobiol 53, 417–424.

Sherman, G., and Visscher, PK. (2002) Honeybee colonies achieve fitness through dancing. Nature 419.6910 (2002): 920–922.

Smarr, B., Burnett, D., Mesri, S., Pister, K. & Kriegsfeld, L. (2015). A Wearable Sensor System with Circadian Rhythm Stability Estimation for Prototyping Biomedical Studies. IEEE Transactions on Affective Computing, PP, 1–1.

Stabentheiner, A., Kovac, H., Mandl, M. and Käfer, H. (2021). Coping with the cold and fighting the heat: thermal homeostasis of a superorganism, the honeybee colony. Journal of Comparative Physiology A, 1–15.

Thomas, C.D. (2010), Climate, climate change and range boundaries. Diversity and Distributions, 16: 488–495.

Tomita, J., Ban, G., & Kume, K. (2017). Genes and neural circuits for sleep of the fruit fly. Neuroscience Research, 118, 82–91.

van Langevelde, F., van Grunsven, R. H., Veenendaal, E. M. and Fijen, T. P. (2017). Artificial night lighting inhibits feeding in moths. Biol Lett 13.

von Frisch, K. (1967). The dance language and orientation of bees. Cambridge, Massachusetts: Belknap Press.

Williams, G. R., Alaux, C., Costa, C., Csaki, T., Doublet, V., Eisenhardt, D., 舰 & Brodschneider, R. (2013). Standard methods for maintaining adult Apis mellifera in cages under in vitro laboratory conditions. Journal of Apicultural Research, 52(1), 1–36

Winston, M. L. (1987). The biology of the honey bee. Cambridge, Massachusetts: Harvard University Press.

Wolf, E., & Zerrahn-Wolf, G. (1935). The dark adaptation of the eye of the honey bee. Journal of General Physiology, 19(2), 229–237.

